# A composition-matching algorithm, MatchIDR, identifies prion-like domains that localize to stress granules

**DOI:** 10.1101/2025.11.24.690216

**Authors:** Sean M. Cascarina, Kacy R. Paul, Larissa L. Ford, Eric D. Ross

**Affiliations:** Department of Biochemistry and Molecular Biology, Colorado State University, Fort Collins, CO 80523, USA

## Abstract

Intrinsically disordered regions (IDRs) play important molecular roles in cells even though they do not adopt a stable structure. Relative to structured regions, IDRs have skewed amino acid compositions favoring polar and charged amino acids. This feature is a major contributor to the biophysical behavior and *in vivo* activity of IDRs, but the relationship between composition and activity depends strongly on which amino acids are enriched within the IDRs. Here, we present a new search algorithm, MatchIDR, that takes as input one or more IDR sequences and finds the nearest compositional matches within a proteome. Using MatchIDR with both artificially designed and native yeast IDRs as query sequences, we successfully identify IDRs from multiple organisms that display predictable enrichment (or non-enrichment) within yeast stress granules. Our results demonstrate that composition-based proteome searches can be an effective strategy for identifying new IDRs with similar *in vivo* activities. MatchIDR is available at https://github.com/RossLabCSU/MatchIDR.

## Introduction

Intrinsically disordered regions (IDRs) are protein segments that do not adopt a stable tertiary structure in the absence of binding partners [1–3]. Although unified by the property of conformational heterogeneity, IDRs can vary substantially with respect to primary sequence, amino acid composition, conformational preferences, biophysical properties, and molecular functions [3]. IDRs tend to evolve faster and tolerate more mutations compared to structured domains [2,4–6]. Consequently, paralogous and orthologous IDRs often diverge considerably with respect to primary sequence. However, IDRs have important biological functions and typically still contain conserved features despite divergence in their primary sequences [7–9].

One important activity recently linked to IDRs is the formation of, or recruitment to, membraneless organelles (MLOs) such as stress granules (SGs), nucleoli, processing bodies, nuclear speckles, and many others [10]. Many MLOs are thought to form via liquid-liquid phase separation of specific proteins or RNAs [10–16]. These key nucleating proteins/RNAs are often referred to as “scaffolds”, whereas proteins and other molecules that are subsequently recruited to MLOs are referred to as “clients” [10,14,17,18]. Current models suggest that IDRs engage in weak, multivalent interactions that can aid in the assembly of MLOs or the recruitment to pre-formed MLOs [19–26]. Importantly, these multivalent interactions are derived from the cumulative effect of key compositional features that are distributed across the length of the IDR. Consequently, for these types of domains, the order of amino acids (i.e., the primary sequence) may be less important than the overall amino acid composition [27].

As proof of principle, we recently screened a large selection of yeast prion-like domains (PrLDs) for the ability to localize to SGs [28]. PrLDs are a specific subcategory of IDRs that have compositional similarity to yeast prion domains; PrLDs are typically enriched in Q and N residues, with secondary enrichment in other uncharged polar and aromatic amino acids [29–32]. SGs are a specific type of MLO that form under acute cellular stress conditions and are enriched in hundreds of different proteins and RNAs [33–37]. Multiple proteins that preferentially localize to SGs contain PrLDs [38,39], and some (but not all) PrLDs are sufficient to drive the protein’s localization to SGs [28,40]. Comparison of the PrLDs that localized to yeast SGs to the PrLDs that did not localize to SGs illuminated the compositional features of PrLDs that favor or disfavor SG localization. These compositional features were sufficient to develop a composition-based algorithm to predict PrLD localization to SGs, as well as design completely artificial PrLDs with or without SG-localization activity, with no consideration for primary sequence [28]. Furthermore, the degree of localization of PrLDs to SGs could be tuned in a stepwise fashion by gradual changes in hydrophobic content: a key feature of PrLDs contributing to SG recruitment [41].

Based on our previous successes in both elucidating and validating the compositional features driving PrLD recruitment to SGs, we reasoned that a composition-based search might identify new IDRs/PrLDs with the ability to localize to SGs [27]. Here we develop a new algorithm, MatchIDR, which scans whole proteomes, identifies, and ranks protein regions that share the highest compositional identity with a user-defined query protein sequence. We show that this approach is effective at identifying sets of domains that predictably do or do not localize to yeast SGs, regardless of the proteome from which the PrLD was identified. These results suggest that a composition-matching strategy is capable of identifying IDRs with specific activities even when their primary sequences differ substantially.

## Results

### The MatchIDR algorithm

Many existing bioinformatic tools search for genes or proteins of interest based on sequence homology to a query sequence [42–49]. To our knowledge, no general-purpose tool has been designed to search for compositional identity regardless of primary sequence identity and/or evolutionary relatedness. To address this need, we developed the MatchIDR algorithm, which can search whole proteomes and identify the region in each protein with the highest compositional identity to an IDR of interest (Fig 1). Compositional identity is based on the total Manhattan distance between the percent compositions for the 20 canonical amino acids (see Materials and Methods). Identified regions are then sorted from highest to lowest compositional identity, revealing the regions of the proteome that are the closest compositional matches to the query sequence.

**Figure 1.**
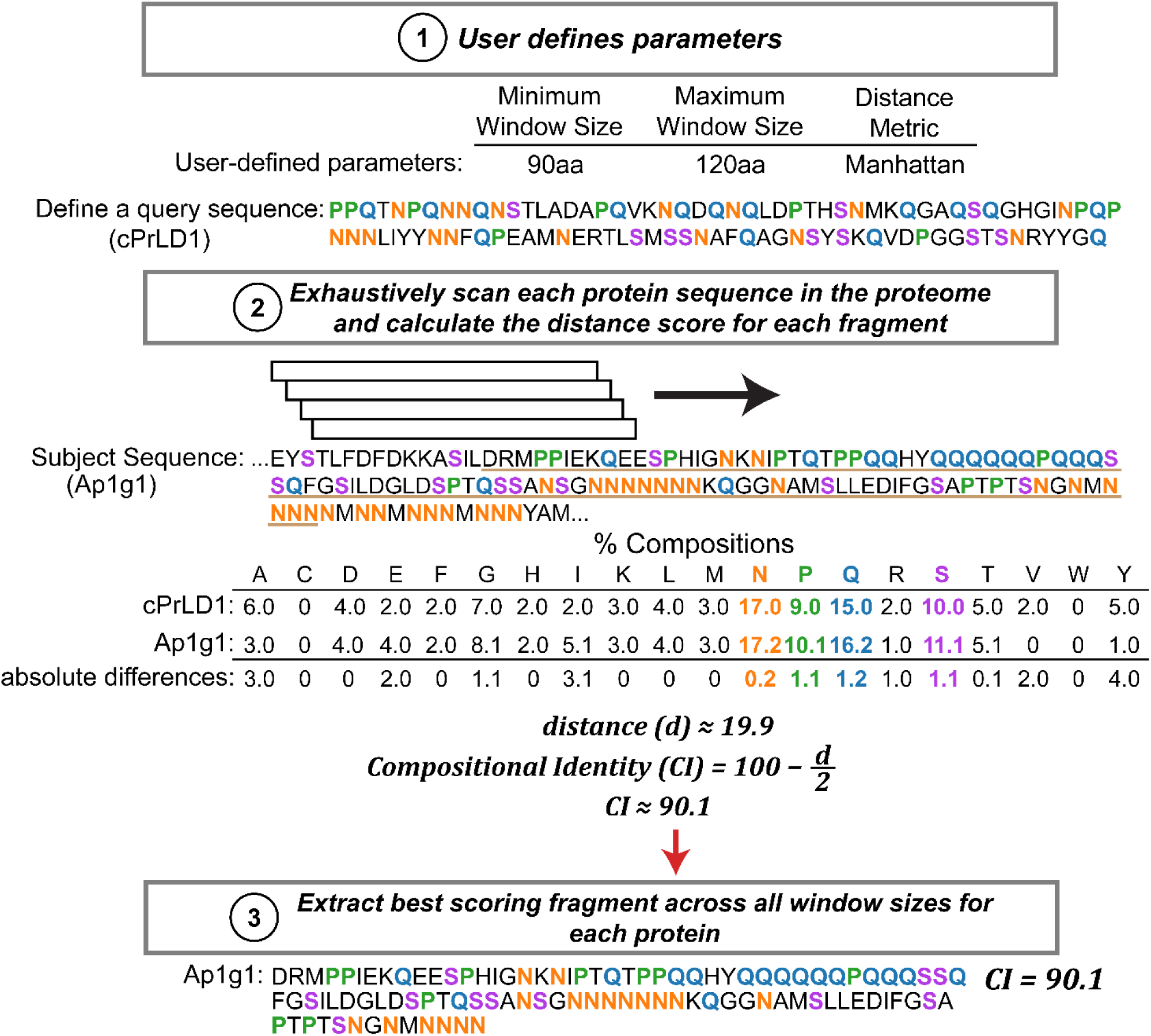
The MatchIDR algorithm. MatchIDR requires a query sequence and a proteome (or any FASTA-formatted file) for composition-matching searches. Additional parameters such as minimum window size, maximum window size, and distance metric can also be specified. MatchIDR scans each protein using the specified window size range and calculates the compositional identity compared to the query sequence. The query sequence in this example is the *in silico*-designed cPrLD1 [28], and the highest scoring fragment corresponds to a region of the slime mold Ap1g1 protein. Prominent compositional features (exceeding 10% of the query or subject sequences) are colored for ease of comparison.

### MatchIDR identifies yeast PrLDs with predictable stress granule localization activity

In prior studies, we elucidated compositional features of PrLDs that favor or disfavor localization to SGs [28,41]. Using these features, we designed synthetic PrLDs (sPrLDs) that localize to SGs during acute heat shock, as well as a complementary set of control PrLDs (cPrLDs) that do not localize to SGs. Since compositional features are sufficient to explain the majority of SG-localization activity for PrLDs, we reasoned that PrLDs with known SG-localization activity could be used as query sequences to identify compositionally similar regions using MatchIDR.

The artificial sPrLD and cPrLD sequences were used as query sequences to identify the top three matches in the yeast proteome for each query PrLD (Table S1; see Materials and Methods for search details). Coding sequences for the composition-matched PrLDs were cloned into a plasmid that tagged them with GFP, then transformed into a strain expressing a cytoplasmic marker (Rpl1b-BFP) and an SG marker (Pab1-mCherry) expressed from endogenous loci. These markers enable quantification of the degree of SG-localization activity encoded by each PrLD, expressed as an “SG enrichment score” where higher values indicate greater enrichment in stress granules [41].

The top matches for the sPrLD and cPrLD sequences exhibit no apparent primary sequence homology but strongly shared compositional features (Fig 2A,B). All three matches for the cPrLD query sequence remain diffuse after acute heat shock even though SGs still form in stressed cells (Fig 2C). This lack of SG localization results in low SG enrichment scores consistent with the original cPrLDs and GFP alone (Fig 2E and Table S2), neither of which are enriched in SGs to a detectable degree. In contrast, the three matches for the sPrLD query sequence form visible foci that colocalize with the Pab1 SG marker after heat shock (Fig 2D). Corresponding SG enrichment scores are consistent with clear SG-localization activity but indicate slight differences in the degree of enrichment within SGs (Fig 2E). All MatchIDR matches remain completely diffuse or mostly diffuse in the absence of heat stress (Fig S1A,B): Ddl1 exhibited occasional pre-stress foci that were not enriched in Pab1 and did not interfere with re-localization to SGs upon heat shock. Therefore, a composition-matching strategy using completely artificial PrLD sequences is sufficient to identify native yeast PrLDs with predictable, stress-responsive SG-localization activity.

**Figure 2.**
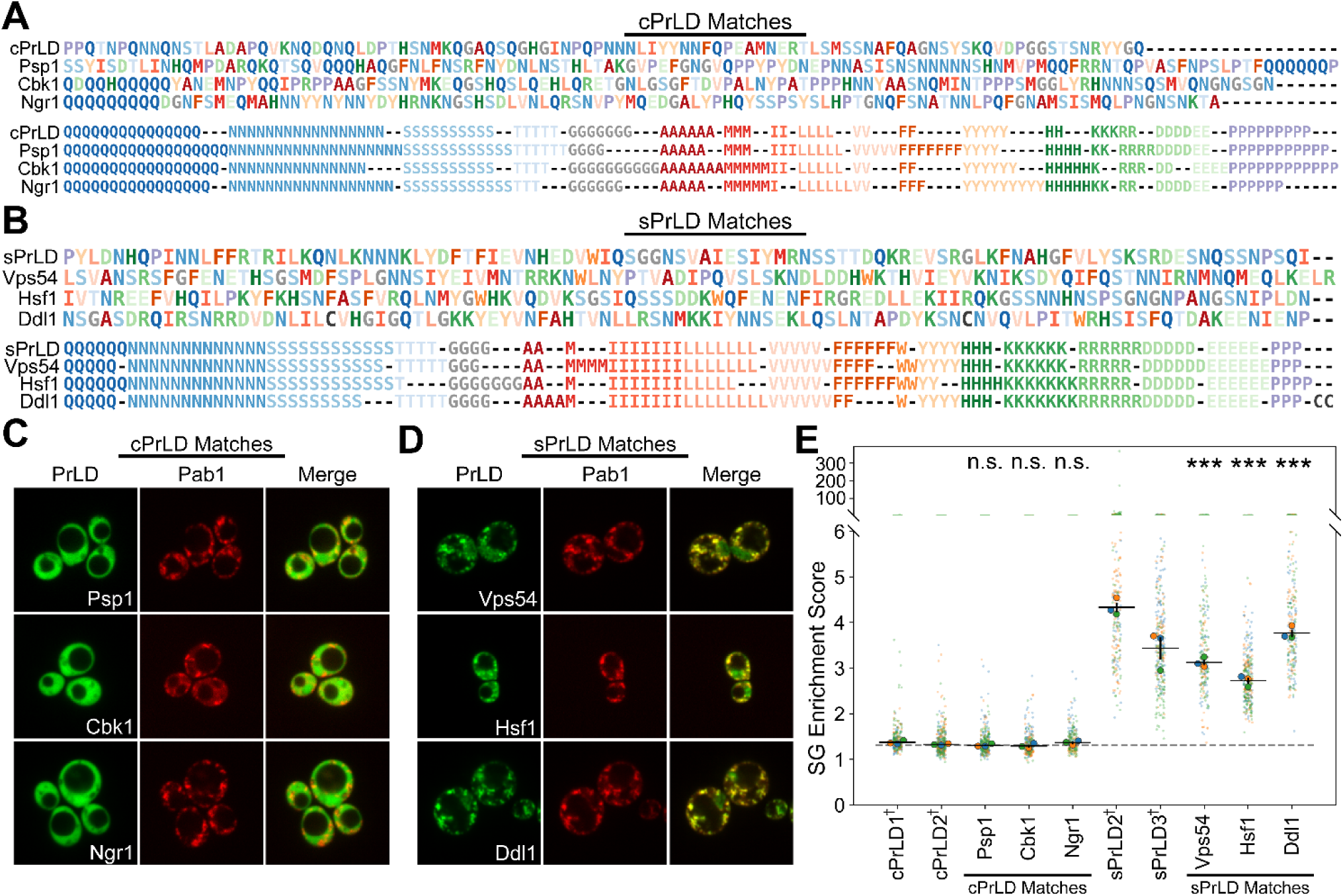
MatchIDR identifies yeast PrLDs with SG-localization activity that matches the sPrLD or cPrLD query sequences. (A) Original sequences (*top*) and composition-clustered sequence representations (*bottom*) of cPrLD and the corresponding top three MatchIDR matches from yeast. (B) Original sequences (*top*) and composition-clustered sequence representations (*bottom*) of sPrLD and the corresponding top three MatchIDR matches from yeast. (C) Representative microscopy images depicting protein localization of the three cPrLD matches and the stress granule marker, Pab1, after 30-minute heat shock at 46°C in yeast. (D) Protein localization of the three sPrLD matches and the stress granule marker, Pab1, after 30-minute heat shock at 46°C in yeast. (E) Quantification of SG enrichment scores for the cPrLD and sPrLD matches. The horizontal dotted line ∼1.3 represents the average of the median SG enrichment scores for GFP alone expressed in the same strain (as reported in [41]) and indicates no detectable enrichment in SGs. All statistical comparisons were performed against cPrLD2 (see Materials and Methods). ^†^Data for cPrLD and sPrLD are derived from our previous publication and are shown here for comparison [41].

### MatchIDR identifies PrLDs from diverse organisms that also exhibit predictable stress granule localization activity in yeast

PrLD localization to SGs is dictated by the overall chemical properties of a PrLD sequence, which is determined largely by amino acid composition. The MatchIDR algorithm requires no organism-specific assumptions: it is equally effective at identifying the best-matching regions regardless of whether the query IDR is derived from the same organism, a different organism, or is entirely synthetic. Therefore, we used MatchIDR with the sPrLD and cPrLD query sequences to identify the top three matches for each sequence from the human and slime mold (*Dictyostelium discoideum*) proteomes using the same search parameters applied to the yeast proteome (Tables S3 and S4).

All matches for the sPrLD or cPrLD sequences share strong compositional identity with each other and with the original query sequence yet very low primary sequence identity in pairwise alignments (Fig 3A,C), indicating that the PrLDs identified by MatchIDR are not simply homologs or paralogs with conserved primary sequences. Furthermore, the PrLD matches exhibit a degree of compositional diversity, with percent-composition ranges for each amino acid typically centered at or near their corresponding query sequence (Fig S2). As observed for yeast, all six cPrLD matches from the human and slime mold proteomes remain diffuse during heat shock (Fig 3B) and exhibit low SG enrichment scores that are consistent with the original cPrLDs and with GFP alone (Fig 3E and Table S2), indicating that none of the cPrLD matches are enriched in SGs. In contrast, all six sPrLD matches from the two proteomes form foci during heat shock (Fig 3D) and have SG enrichment scores that are, on average, higher than scores for cPrLD and GFP alone (Fig 3E and Table S2). However, the degree of SG localization is not correlated with the magnitude of compositional identity of each match compared to the corresponding query sequence (Fig S3). Most human and slime mold PrLDs were completely diffuse prior to heat shock (Fig S1A,B). While the slime mold matches with the sPrLD query protein formed non-SG foci in a subset of cells prior to stress (Fig S1B), all three PrLDs robustly re-localized to SGs after heat shock (Fig 3D). Collectively, these results suggest that the composition-matching strategy is excellent at identifying candidates with or without the intended activity but does not precisely predict the level of SG-localization activity, as deviations in composition could diminish, enhance, or have little effect on activity.

**Figure 3.**
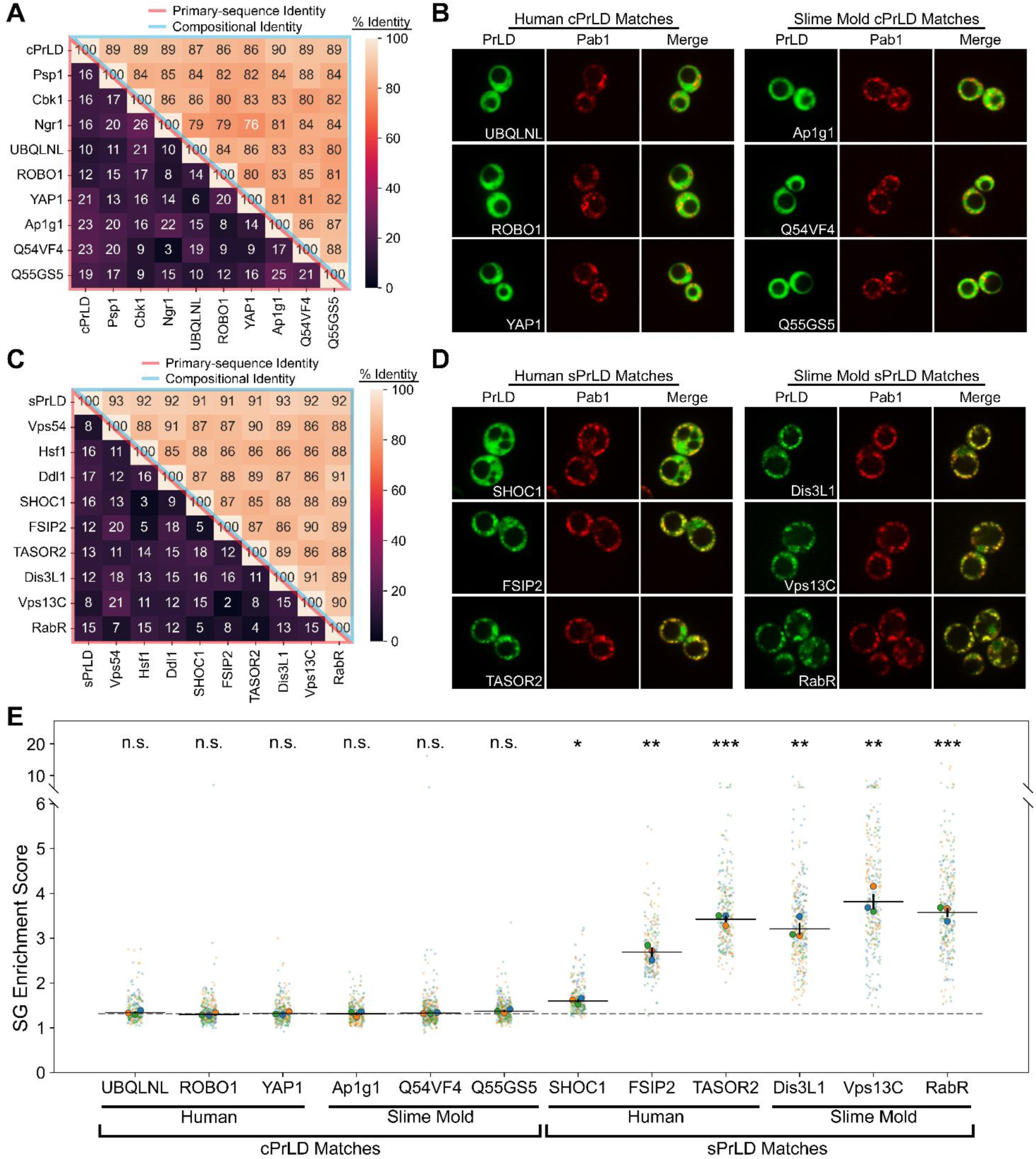
SG-localization activity of composition-matched PrLDs from eukaryotic organisms. (A) Heatmap depicting the primary-sequence identity (*lower-left half*) and the compositional identity (*upper-right half*) for the top three PrLD matches for the cPrLD query sequence from yeast, humans, and slime mold. (B) Representative images displaying the protein localization of each cPrLD match from humans and slime mold with Pab1-positive SGs that form in response to heat shock. (C) Heatmap depicting the primary-sequence identity (*lower-left half*) and the compositional identity (*upper-right half*) for the top three PrLD matches for the sPrLD query sequence from yeast, humans, and slime mold. (D) Representative images displaying the colocalization of each sPrLD match from humans and slime mold with Pab1-positive SGs that form in response to heat shock. (E) Quantification of SG enrichment scores for each PrLD. The horizontal dotted line represents the average of the median SG enrichment scores for GFP alone, as reported in [41]. All statistical comparisons were performed against cPrLD2 (see Materials and Methods).

### Native yeast PrLDs are effective query sequences for identifying human PrLDs with SG-localization activity

In a prior study, we estimated SG propensity scores for each amino acid and used these scores to develop a composition-based SG prediction algorithm for PrLDs, as well as design the sPrLD and cPrLD sequences [28]. These sPrLDs and cPrLDs were artificially engineered for high SG enrichment and no SG enrichment, respectively, making them ideal starting points to test the effectiveness of the composition-matching strategy employed by MatchIDR. To examine whether this strategy would also work when native yeast PrLDs are used as query sequences, we chose three native yeast PrLDs with known SG-localization activity and three native yeast PrLDs that do not localize to SGs [28]. Importantly, to determine if the composition-matching strategy could be effective even in cases where the compositional features driving activity are not fully understood, we specifically chose PrLDs that exhibited SG-enrichment activity that was poorly predicted by our SG prediction algorithm (Fig S4; [28]). For each PrLD, a MatchIDR search was performed for the human proteome using the same parameters described for the sPrLD/cPrLD searches (Table S5). In each case, the top five human PrLD candidates were evaluated using our SG scoring algorithm [28]. The single candidate with an SG score most closely matching the SG score of the yeast query PrLD was tested for SG-localization activity.

As with the sPrLD and cPrLD searches, the single best match exhibited high compositional identity but low primary sequence identity compared to the corresponding query sequence (Fig 4A). In contrast, comparisons across non-similar PrLDs yielded lower compositional and primary sequence identities, reflecting the unique compositional profile of each native yeast PrLD used as query sequences. Among the set of human PrLDs expected to remain diffuse during heat shock, two out of the three human PrLD matches fail to form foci (Fig 4B) and have correspondingly low SG enrichment scores consistent with no detectable SG localization (Fig 4D and Table S2). Of the six human PrLDs, only ARMCX2 showed substantial foci formation prior to stress (Fig S1C,D), potentially contributing to its unexpected ability to localize to SGs. In contrast, all three human PrLD matches that were expected to localize to SGs form foci (Fig 4C) and have high SG enrichment scores (Fig 4D and Table S2), consistent with recruitment of these PrLDs to SGs. However, the degree of SG enrichment of the human PrLDs differed from that of the corresponding yeast query sequence, and differences in SG enrichment were not correlated with compositional identity (Fig S5). Therefore, as with the sPrLD and cPrLD searches, MatchIDR run with native yeast PrLDs is effective at identifying composition-matched regions with or without SG-localization activity but does not predict the degree of SG enrichment. This highlights that MatchIDR can retrieve sequences with similar activity from whole-proteome searches even in cases where the compositional requirements for the activity are poorly understood.

**Figure 4.**
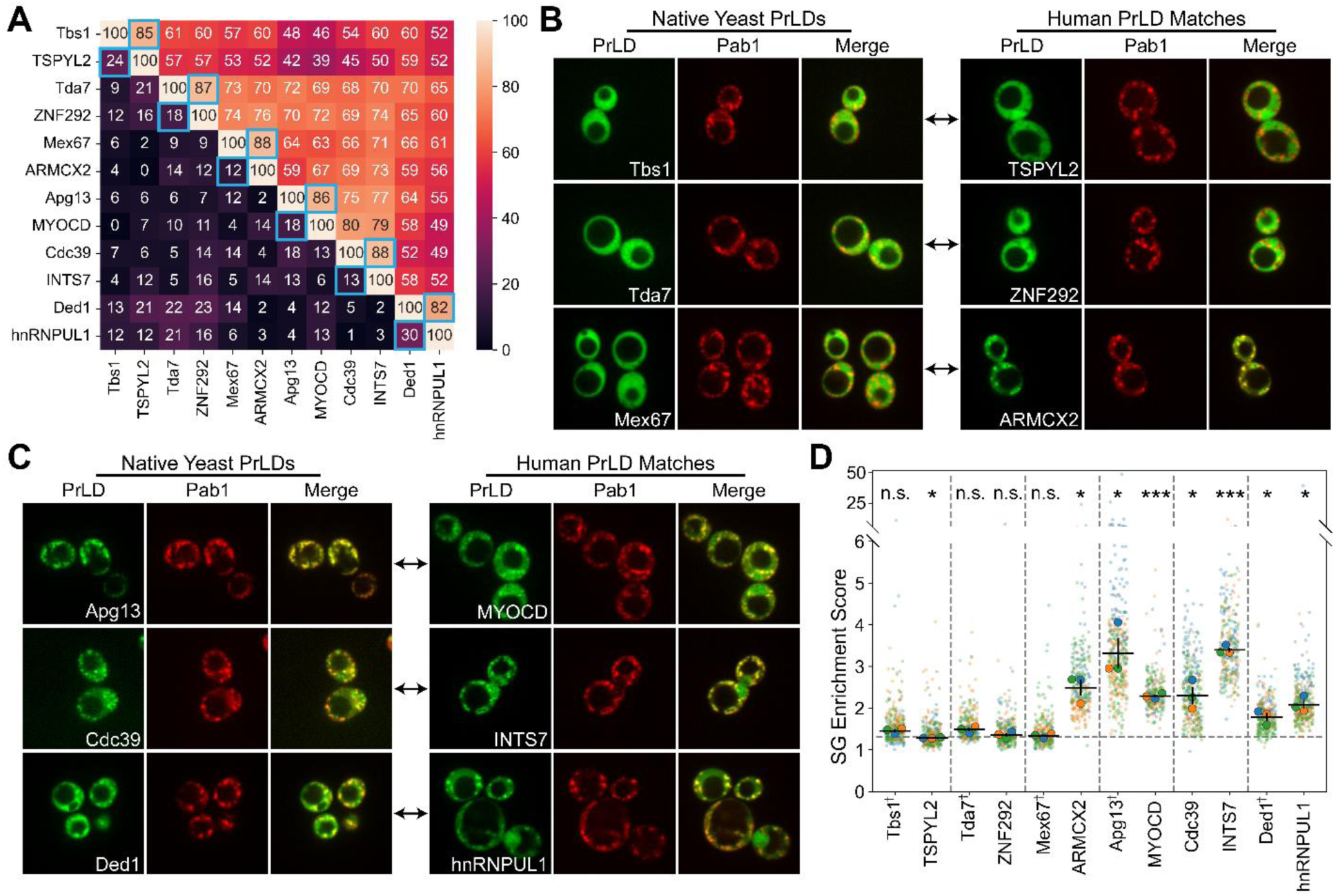
SG-localization activity of composition-matched human PrLDs for the native yeast query PrLDs. (A) Heatmap depicting the primary-sequence identity (*lower-left half*) and the compositional identity (*upper-right half*) for the single best human PrLD matches for each yeast query sequence. Each yeast query PrLD appears before its paired human PrLD match, and these pairs are highlighted in adjacent blue boxes. (B) Representative microscopy images depicting protein localization of the three native yeast PrLDs that do not localize to SGs (*left images*) and their corresponding top human PrLD matches (*right images*) after 30-minute heat shock in yeast. SGs formed during heat shock in panels B and C are indicated by Pab1 foci. (C) Representative microscopy images depicting protein localization of the three native yeast PrLDs that localize to SGs (*left images*) and their corresponding top human PrLD matches (*right images*) after 30-minute heat shock in yeast. (D) Quantification of SG enrichment scores for each PrLD. The horizontal dotted line represents the average of the median SG enrichment scores for GFP alone, as reported in [41]. All statistical comparisons were performed against cPrLD2 (see Methods). ^†^PrLDs with one replicate (blue) derived from our previous study [41].

## Discussion

Protein sequences encoding well-folded domains are often evolutionarily constrained and highly specific for the structures and activities that they encode. This tight relationship between amino acid sequence and protein activity limits the number and magnitude of sequence changes that can be tolerated by a sequence while still maintaining its activity. IDRs tend to be less constrained in this regard, often exhibiting higher rates of evolutionary divergence and large deviations in primary sequence. The overall amino acid composition of IDRs is often preserved despite changes in primary sequence [7–9,50–52]. This suggests that, for some IDRs, their amino acid composition is a stronger determinant of activity than their primary sequence, which we refer to as a composition-driven protein activity [27].

Based on this hypothesis, we developed a computational tool, MatchIDR, to search for the nearest compositional match to a query sequence without regard for primary sequence. Although MatchIDR can perform cross-organism searches, the goal of this search tool is not necessarily to identify homologous domains: rather, it is to identify domains with the greatest similarity in terms of compositional features. For query proteins or domains with composition-driven activities, top-scoring candidates are likely to encode similar activities through the features that they share with the query proteins.

As proof of principle, we employed MatchIDR to examine the efficacy of a composition-based proteome search method for identifying SG-localizing PrLDs. Multiple lines of evidence support a composition-driven activity model for SGs [27]. PrLDs that localize to SGs share compositional biases but differ substantially in terms of primary sequence [28]. Scrambled PrLD sequences, which have identical compositions but distinct primary sequences, still localize to SGs [28]. Compositional features alone can make accurate predictions of SG-localization behavior of PrLDs and are sufficient to design artificial sequences with predictable SG localization [28]. Rational changes to compositional features – without consideration of primary sequence – lead to predictable, stepwise changes in SG localization [41].

Consistent with a composition-driven model of PrLD localization to SGs, MatchIDR was remarkably successful at identifying PrLDs with SG-localization activity that mirrors their query proteins. Query proteins that localize to SGs identified new PrLDs that also encoded SG-localization activity, whereas query proteins that do not localize to SGs yielded new PrLDs that also do not localize to SGs. This approach was effective regardless of the organism that was searched and despite low primary sequence homologies among the matches, suggesting that the effect was driven largely by the chemical composition of the identified domains rather than conserved primary sequence motifs or yeast-specific binding partners. However, it is important to note that SG-localization activity may have different requirements in different organisms. For consistency, we tested all domains in our yeast model organism, where the original compositional features of SG-localization activity were initially elucidated. While many human and slime mold PrLDs localized to SGs in yeast, these same PrLDs may not localize to SGs in their native host organisms due to different compositional or sequence requirements in that environment.

Collectively, these results provide further evidence that PrLD localization to SGs is driven by amino acid composition. When applied to these types of proteins and activities, MatchIDR is a powerful tool to uncover additional protein regions with shared compositional features and encoded activities.

## Materials and Methods

### Yeast Strains, Cloning, and Growth Conditions

Experiments testing the localization of PrLDs to SGs were performed as previously described [41]. Briefly, each PrLD identified by MatchIDR was cloned into a vector to express the PrLD with an N-terminal GFP tag from the *ADH1* promoter. For the native yeast PrLDs used as query sequences for MatchIDR searches, we utilized previously constructed vectors expressing the PrLD with an N-terminal GFP tag from the *SUP35* promoter [28]: in our recent study, the choice of promoter did not appreciably affect PrLD enrichment in SGs [41]. These vectors were transformed into a strain expressing the cytoplasmic marker (Rpl1b-BFP) and SG marker (Pab1-mCherry) from their endogenous loci. Cells were grown overnight in SC-Leu media (synthetic complete media lacking leucine) then diluted into fresh media, grown to mid-log phase, and subjected to a 30-minute heat shock at 46°C immediately prior to imaging. At least three experimental replicates were performed for each PrLD, with a minimum of 30 quantifiable cells per replicate.

### Quantification of SG Enrichment Scores

All microscopy images were collected on an Olympus (IX83) inverted spinning-disk confocal microscope using a 100x objective. SG enrichment scores were calculated from microscopy images using our previously developed pipeline [41]. Yeast cells were initially identified using YeastSpotter [53]. Prior to quantification, cells were screened using a variety of criteria to ensure quality single-cell data (see [41] for a full description), with one minor screening change: cells with little or no PrLD-GFP expression were defined as those with a median cytoplasmic GFP intensity <1000 and a 90^th^ percentile (rather than the median) GFP intensity in SGs <1000. The cytoplasm of each cell was defined as pixels with >0.8x the median intensity for all pixels in the cell for the Rpl1b-BFP cytoplasmic marker. SGs were classified as regions with pixel intensities >2x the median intensity in the cell for the Pab1-mCherry SG marker. SG enrichment for the PrLD-GFP was calculated for each cell as the 90^th^ percentile of GFP intensities in the SGs divided by the median GFP intensity in non-SG cytoplasmic regions [41]. For all statistical comparisons, the median for each replicate was calculated from log_2_-transformed, single-cell SG enrichment scores. Medians were then averaged across replicates to yield a mean SG enrichment score. The mean SG enrichment score for each PrLD was compared to the mean SG enrichment score of cPrLD2 using a two-sided Welch’s *t* test. Statistical significance indicators (p>0.05, “n.s.”; p<0.05, *; p<0.01, **; p<0.001, ***) in all figures summarize raw *p*-values from these comparisons (Table S2).

### Calculation of Primary Sequence Identity and Compositional Identity

PrLDs were compared to the original query sequences and to each other in a pairwise fashion with respect to both primary sequence and composition. Primary sequence identity was extracted from pairwise alignments using the EMBOSS Needle program [54]. Compositional identity (*CI*) was calculated according to the following equation:

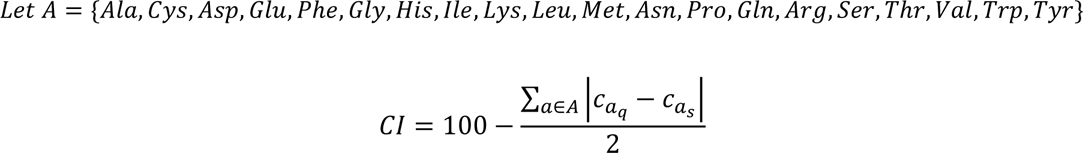

where *A* represents the set of canonical amino acids and *c* represents the percent composition of amino acid *a* in the query *q* or subject *s* sequences. The sum term in the numerator represents the total Manhattan distance for the percent compositions for the 20 canonical amino acids in a subject sequence compared to the query sequence.

### MatchIDR Search Parameters for sPrLD, cPrLD, and Native Yeast PrLD Query Sequences

The minimum length of PrLDs capable of driving localization to SGs has not been empirically determined. We used three main observations from our previous study [28] to determine the window size range used for MatchIDR searches: 1) multiple native yeast PrLDs that are 81 amino acids in length can localize to SGs; 2) two sPrLDs that are 100 amino acids in length are strongly enriched in SGs; and 3) the median length of native yeast PrLDs that localize to SGs was ∼122 amino acids. Therefore, we chose a narrow window size range of 90-120 amino acids (inclusive) for MatchIDR searches to identify candidate PrLDs to test in SG localization experiments. All MatchIDR searches were performed using the default Manhattan distance metric. Proteomes involved in searches include the yeast (*Saccharomyces cerevisiae*), slime mold (*Dictyostelium discoideum*), and human (*Homo sapiens*) proteomes (UniProt IDs UP000002311_559292, UP000002195_44689, and UP000005640_9606, respectively). The MatchIDR Python script is available at https://github.com/RossLabCSU/MatchIDR.

## Notes

### Competing Interest Statement

The authors have declared no competing interest.

### Funding

- National Institute of General Medical Sciences: R35GM130352

## References

1. Lemke EA, Babu MM, Kriwacki RW, Mittag T, Pappu R V., Wright PE, et al. Intrinsic disorder: A term to define the specific physicochemical characteristic of protein conformational heterogeneity. Mol Cell. 2024;84: 1188–1190. doi:10.1016/j.molcel.2024.02.024

2. van der Lee R, Buljan M, Lang B, Weatheritt RJ, Daughdrill GW, Dunker AK, et al. Classification of Intrinsically Disordered Regions and Proteins. Chem Rev. 2014;114: 6589–6631. doi:10.1021/cr400525m

3. Holehouse AS, Kragelund BB. The molecular basis for cellular function of intrinsically disordered protein regions. Nat Rev Mol Cell Biol. 2024;25: 187–211. doi:10.1038/s41580-023-00673-0

4. Brown CJ, Johnson AK, Dunker AK, Daughdrill GW. Evolution and disorder. Curr Opin Struct Biol. 2011;21: 441–446. doi:10.1016/j.sbi.2011.02.005

5. Bellay J, Han S, Michaut M, Kim T, Costanzo M, Andrews BJ, et al. Bringing order to protein disorder through comparative genomics and genetic interactions. Genome Biol. 2011;12: R14. doi:10.1186/gb-2011-12-2-r14

6. Brown CJ, Takayama S, Campen AM, Vise P, Marshall TW, Oldfield CJ, et al. Evolutionary Rate Heterogeneity in Proteins with Long Disordered Regions. J Mol Evol. 2002;55: 104–110. doi:10.1007/s00239-001-2309-6

7. Zarin T, Strome B, Nguyen Ba AN, Alberti S, Forman-Kay JD, Moses AM. Proteome-wide signatures of function in highly diverged intrinsically disordered regions. Elife. 2019;8: e46883. doi:10.7554/eLife.46883

8. Zarin T, Strome B, Peng G, Pritišanac I, Forman-Kay JD, Moses AM. Identifying molecular features that are associated with biological function of intrinsically disordered protein regions. Elife. 2021;10: e60220. doi:10.7554/eLife.60220

9. Moesa HA, Wakabayashi S, Nakai K, Patil A. Chemical composition is maintained in poorly conserved intrinsically disordered regions and suggests a means for their classification. Mol Biosyst. 2012;8: 3262–3273. doi:10.1039/c2mb25202c

10. Banani SF, Lee HO, Hyman AA, Rosen MK. Biomolecular condensates: organizers of cellular biochemistry. Nature Reviews Molecular Cell Biology. 2017. pp. 285–298. doi:10.1038/nrm.2017.7

11. Guillén-Boixet J, Kopach A, Holehouse AS, Wittmann S, Jahnel M, Schlüßler R, et al. RNA-Induced Conformational Switching and Clustering of G3BP Drive Stress Granule Assembly by Condensation. Cell. 2020;181: 346–361. doi:10.1016/j.cell.2020.03.049

12. Yang P, Mathieu C, Kolaitis RM, Zhang P, Messing J, Yurtsever U, et al. G3BP1 Is a Tunable Switch that Triggers Phase Separation to Assemble Stress Granules. Cell. 2020;181: 325–345. doi:10.1016/j.cell.2020.03.046

13. Sanders DW, Kedersha N, Lee DSW, Strom AR, Drake V, Riback JA, et al. Competing Protein-RNA Interaction Networks Control Multiphase Intracellular Organization. Cell. 2020;181: 306–324. doi:10.1016/j.cell.2020.03.050

14. Xing W, Muhlrad D, Parker R, Rosen MK. A quantitative inventory of yeast P body proteins reveals principles of composition and specificity. Elife. 2020;9: e56525. doi:10.7554/eLife.56525

15. Begovich K, Wilhelm JE. An In Vitro Assembly System Identifies Roles for RNA Nucleation and ATP in Yeast Stress Granule Formation. Mol Cell. 2020;79: 991–1007. doi:10.1016/j.molcel.2020.07.017

16. Freibaum BD, Messing J, Yang P, Kim HJ, Taylor JP. High-fidelity reconstitution of stress granules and nucleoli in mammalian cellular lysate. Journal of Cell Biology. 2021;220: e202009079. doi:10.1083/jcb.202009079

17. Ditlev JA, Case LB, Rosen MK. Who’s In and Who’s Out—Compositional Control of Biomolecular Condensates. J Mol Biol. 2018;430: 4666–4684. doi:10.1016/j.jmb.2018.08.003

18. Banani SF, Rice AM, Peeples WB, Lin Y, Jain S, Parker R, et al. Compositional Control of Phase-Separated Cellular Bodies. Cell. 2016;166: 651–663. doi:10.1016/j.cell.2016.06.010

19. Protter DSW, Rao BS, Van Treeck B, Lin Y, Mizoue L, Rosen MK, et al. Intrinsically Disordered Regions Can Contribute Promiscuous Interactions to RNP Granule Assembly. Cell Rep. 2018;22: 1401–1412. doi:10.1016/j.celrep.2018.01.036

20. Nott TJ, Petsalaki E, Farber P, Jervis D, Fussner E, Plochowietz A, et al. Phase Transition of a Disordered Nuage Protein Generates Environmentally Responsive Membraneless Organelles. Mol Cell. 2015;57: 936–947. doi:10.1016/j.molcel.2015.01.013

21. Elbaum-Garfinkle S, Kim Y, Szczepaniak K, Chen CC-H, Eckmann CR, Myong S, et al. The disordered P granule protein LAF-1 drives phase separation into droplets with tunable viscosity and dynamics. Proc Natl Acad Sci U S A. 2015;112: 7189–7194. doi:10.1073/pnas.1504822112

22. Molliex A, Temirov J, Lee J, Coughlin M, Kanagaraj AP, Kim HJ, et al. Phase Separation by Low Complexity Domains Promotes Stress Granule Assembly and Drives Pathological Fibrillization. Cell. 2015;163: 123–133. doi:10.1016/j.cell.2015.09.015

23. Martin EW, Holehouse AS, Peran I, Farag M, Incicco JJ, Bremer A, et al. Valence and patterning of aromatic residues determine the phase behavior of prion-like domains. Science (1979). 2020;367: 694–699. doi:10.1126/science.aaw8653

24. Bremer A, Farag M, Borcherds WM, Peran I, Martin EW, Pappu R V., et al. Deciphering how naturally occurring sequence features impact the phase behaviours of disordered prion-like domains. Nat Chem. 2022;14: 196–207. doi:10.1038/s41557-021-00840-w

25. Pak CW, Kosno M, Holehouse AS, Padrick SB, Mittal A, Ali R, et al. Sequence Determinants of Intracellular Phase Separation by Complex Coacervation of a Disordered Protein. Mol Cell. 2016;63: 72–85. doi:10.1016/j.molcel.2016.05.042

26. Choi J-M, Holehouse AS, Pappu R V. Physical Principles Underlying the Complex Biology of Intracellular Phase Transitions. Annu Rev Biophys. 2020;49: 107–133. doi:10.1146/annurev-biophys-121219-081629

27. Cascarina SM, Ross ED. Protein activities driven by amino acid composition. Journal of Biological Chemistry. 2025; 110640. doi:10.1016/j.jbc.2025.110640

28. Boncella AE, Shattuck JE, Cascarina SM, Paul KR, Baer MH, Fomicheva A, et al. Composition-based prediction and rational manipulation of prion-like domain recruitment to stress granules. Proc Natl Acad Sci U S A. 2020;117: 5826–5835. doi:10.1073/pnas.1912723117

29. Cascarina SM, Ross ED. Yeast prions and human prion-like proteins: sequence features and prediction methods. Cellular and Molecular Life Sciences. 2014/01/07. 2014;71: 2047–2063. doi:10.1007/s00018-013-1543-6

30. Alberti S, Halfmann R, King O, Kapila A, Lindquist S. A Systematic Survey Identifies Prions and Illuminates Sequence Features of Prionogenic Proteins. Cell. 2009;137: 146–158. doi:10.1016/j.cell.2009.02.044

31. Michelitsch MD, Weissman JS. A census of glutamine/asparagine-rich regions: Implications for their conserved function and the prediction of novel prions. Proc Natl Acad Sci U S A. 2000;97: 11910–11915. doi:10.1073/pnas.97.22.11910

32. Harrison PM, Gerstein M. A method to assess compositional bias in biological sequences and its application to prion-like glutamine/asparagine-rich domains in eukaryotic proteomes. Genome Biol. 2003;4: R40. doi:10.1186/gb-2003-4-6-r40

33. Jain S, Wheeler JR, Walters RW, Agrawal A, Barsic A, Parker R. ATPase-Modulated Stress Granules Contain a Diverse Proteome and Substructure. Cell. 2016;164: 487–498. doi:10.1016/j.cell.2015.12.038

34. Khong A, Matheny T, Jain S, Mitchell SF, Wheeler JR, Parker R. The Stress Granule Transcriptome Reveals Principles of mRNA Accumulation in Stress Granules. Mol Cell. 2017;68: 808–820. doi:10.1016/j.molcel.2017.10.015

35. Markmiller S, Soltanieh S, Server KL, Mak R, Jin W, Fang MY, et al. Context-Dependent and Disease-Specific Diversity in Protein Interactions within Stress Granules. Cell. 2018;172: 590–604.e13. doi:10.1016/j.cell.2017.12.032

36. Marmor-Kollet H, Siany A, Kedersha N, Knafo N, Rivkin N, Danino YM, et al. Spatiotemporal Proteomic Analysis of Stress Granule Disassembly Using APEX Reveals Regulation by SUMOylation and Links to ALS Pathogenesis. Mol Cell. 2020;80: 876–891.e6. doi:10.1016/j.molcel.2020.10.032

37. Protter DSW, Parker R. Principles and Properties of Stress Granules. Trends in Cell Biology. 2016. pp. 668–679. doi:10.1016/j.tcb.2016.05.004

38. Harrison AF, Shorter J. RNA-binding proteins with prion-like domains in health and disease. Biochemical Journal. 2017;474: 1417–1438. doi:10.1042/BCJ20160499

39. Li YR, King OD, Shorter J, Gitler AD. Stress granules as crucibles of ALS pathogenesis. Journal of Cell Biology. 2013/05/01. 2013;201: 361–372. doi:10.1083/jcb.201302044

40. Shattuck JE, Paul KR, Cascarina SM, Ross ED. The prion-like protein kinase Sky1 is required for efficient stress granule disassembly. Nat Commun. 2019. doi:10.1038/s41467-019-11550-w

41. Baer MH, Cascarina SM, Paul KR, Ross ED. Rational Tuning of the Concentration-independent Enrichment of Prion-like Domains in Stress Granules. J Mol Biol. 2024;436: 168703. doi:10.1016/j.jmb.2024.168703

42. Altschul SF, Gish W, Miller W, Myers EW, Lipman DJ. Basic local alignment search tool. J Mol Biol. 1990;215: 403–410. doi:10.1016/S0022-2836(05)80360-2

43. Pearson WR, Lipman DJ. Improved tools for biological sequence comparison. Proc Natl Acad Sci U S A. 1988;85: 2444–2448. doi:10.1073/pnas.85.8.2444

44. Steinegger M, Söding J. MMseqs2 enables sensitive protein sequence searching for the analysis of massive data sets. Nat Biotechnol. 2017;35: 1026–1028. doi:10.1038/nbt.3988

45. Li J, Wang Z, Fan X, Yao R, Zhang G, Fan R, et al. Rapid multiple protein sequence search by parallel and heterogeneous computation. Bioinformatics. 2024;40. doi:10.1093/bioinformatics/btae151

46. Buchfink B, Xie C, Huson DH. Fast and sensitive protein alignment using DIAMOND. Nat Methods. 2014;12: 59–60. doi:10.1038/nmeth.3176

47. Edgar RC. Search and clustering orders of magnitude faster than BLAST. Bioinformatics. 2010;26: 2460–2461. doi:10.1093/bioinformatics/btq461

48. Kiełbasa SM, Wan R, Sato K, Horton P, Frith MC. Adaptive seeds tame genomic sequence comparison. Genome Res. 2011;21: 487–493. doi:10.1101/gr.113985.110

49. Eddy SR. Accelerated Profile HMM Searches. PLoS Comput Biol. 2011;7: e1002195. doi:10.1371/journal.pcbi.1002195

50. Adiji OA, McConnell BS, Parker MW. The origin recognition complex requires chromatin tethering by a hypervariable intrinsically disordered region that is functionally conserved from sponge to man. Nucleic Acids Res. 2024;52: 4344–4360. doi:10.1093/nar/gkae122

51. Santoso A, Chien P, Osherovich LZ, Weissman JS. Molecular Basis of a Yeast Prion Species Barrier. Cell. 2000;100: 277–288. doi:10.1016/S0092-8674(00)81565-2

52. Chernoff YO, Galkin AP, Lewitin E, Chernova TA, Newnam GP, Belenkiy SM. Evolutionary conservation of prion-forming abilities of the yeast Sup35 protein. Mol Microbiol. 2000;35: 865–876. doi:10.1046/j.1365-2958.2000.01761.x

53. Lu AX, Zarin T, Hsu IS, Moses AM. YeastSpotter: accurate and parameter-free web segmentation for microscopy images of yeast cells. Bioinformatics. 2019;35: 4525–4527. doi:10.1093/bioinformatics/btz402

54. Rice P, Longden I, Bleasby A. EMBOSS: The European Molecular Biology Open Software Suite. Trends in Genetics. 2000;16: 276–277. doi:10.1016/S0168-9525(00)02024-2

