## Supplemental_Materials for "A composition-matching algorithm, MatchIDR, identifies prion-like domains that localize to stress granules"

### Supplementary Figures

**Fig S1. Representative images of yeast expressing MatchIDR-identified PrLDs at the standard 30°C growth temperature.** Most PrLDs exhibited diffuse cytoplasmic localization at 30°C. For PrLDs with a small but reproducible subset of cells forming foci at 30°C (Ddl1, Dis3L1, Vps13C, RabR, and ARMCX2), representative images were selected to show at least one cell with a focus but do not reflect the frequency of foci-forming cells among the population.

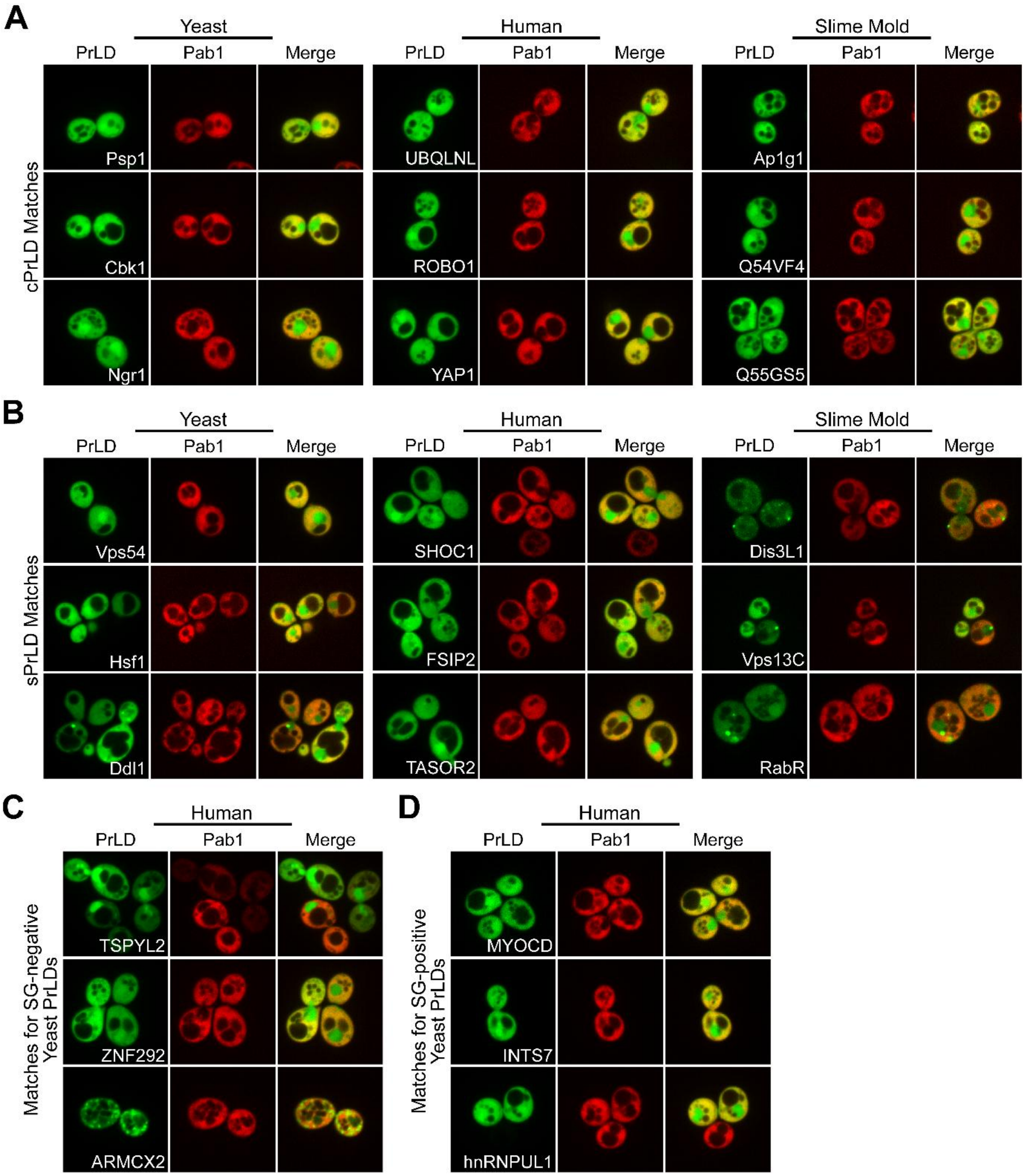

**Fig S2. Amino acid composition of sPrLD matches and cPrLD matches.** Percent composition for each amino acid was calculated for MatchIDR matches from yeast, humans, and slime mold and grouped based on the original query protein (sPrLD matches in blue, cPrLD matches in red). For comparison, the composition values for sPrLD and cPrLD are represented as black stars.

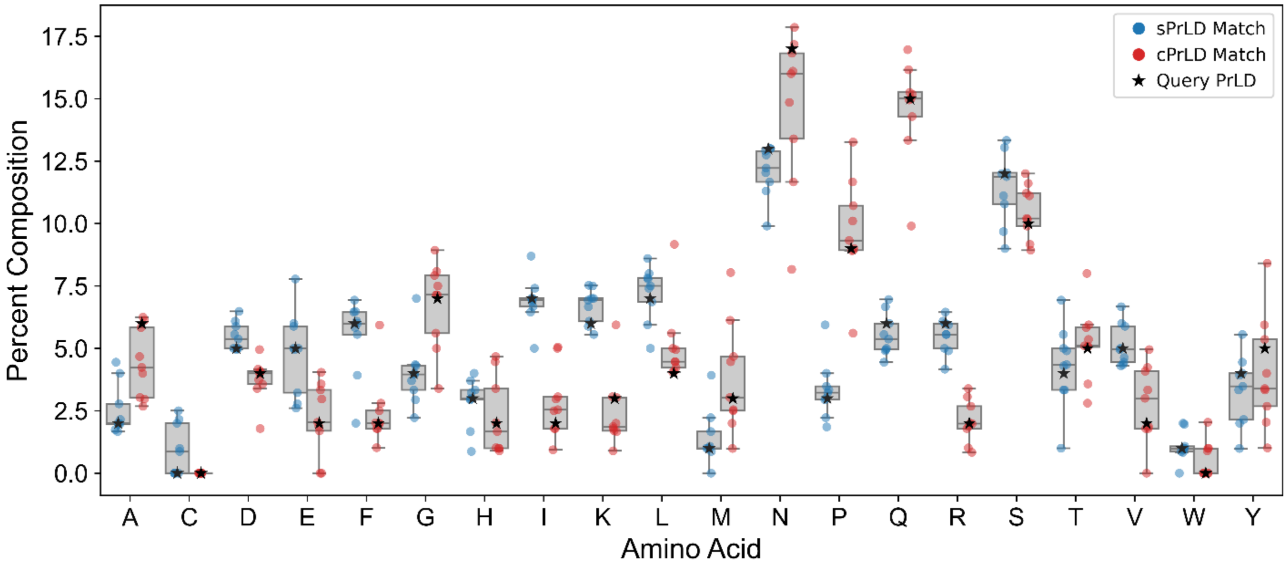

**Fig S3. SG enrichment score versus compositional identity for sPrLD and cPrLD matches.** MatchIDR matches for sPrLD (blue) and cPrLD (red) from yeast, humans, and slime mold are plotted with respect to mean SG enrichment score and compositional identity relative to their corresponding reference sequences.

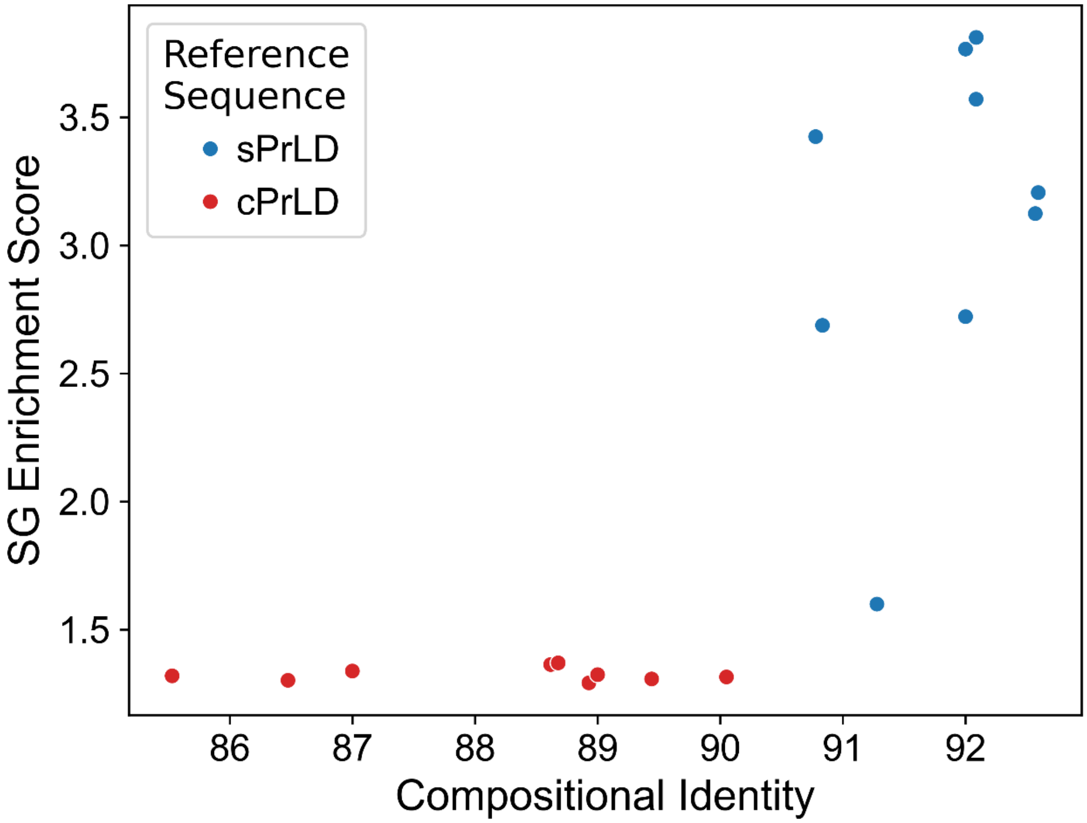

**Fig S4. Stress granule propensity scores for native yeast PrLDs with previously characterized SG-localization activity.** Predicted SG propensities for a set of native yeast PrLDs with SG enrichment (“SG-positive PrLDs”) or no detectable SG enrichment (“SG-negative PrLDs”) were reported in our prior study (Boncella and Shattuck *et al.*, 2020). The contribution of each amino acid to SG localization was estimated based on their enrichment or depletion in the SG-positive PrLD sequences relative to the SG-negative PrLD sequences. Predicted SG propensity scores are the average estimated contribution of each residue across the PrLD sequence. The six native yeast PrLDs used as query sequences in MatchIDR searches (labeled proteins; this study) represent the most extreme inconsistencies between predicted SG propensity and observed SG-localization activity.

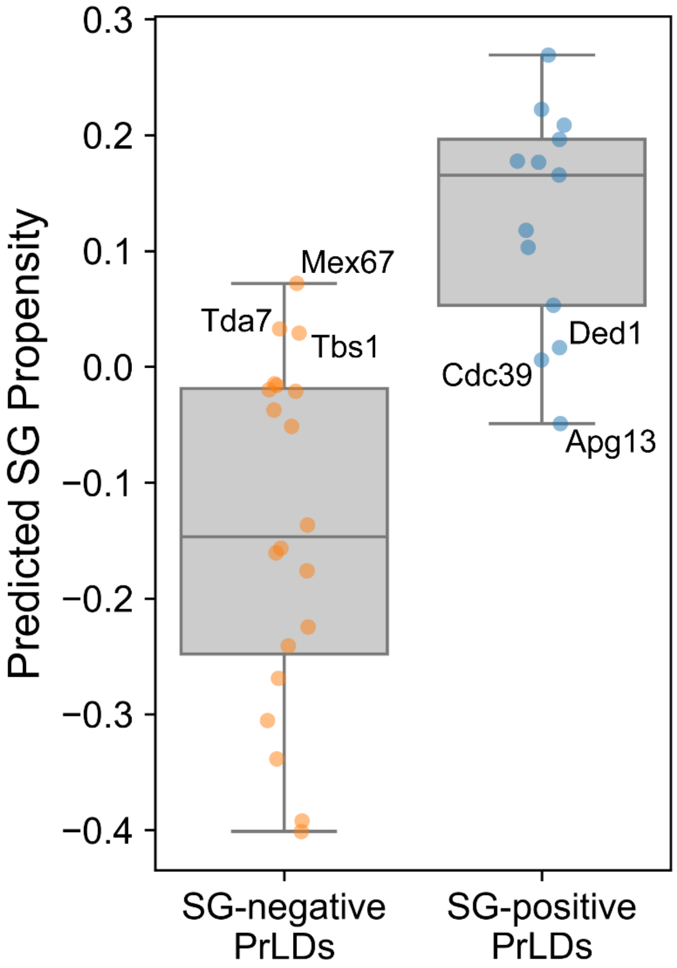

**Fig S5. SG enrichment score versus compositional identity for human PrLDs identified as top matches using native yeast PrLDs query sequences.** Each human MatchIDR match was compared to its corresponding yeast PrLD query sequence with respect to both SG enrichment score and compositional identity. Values on the *y*-axis represent the difference in mean SG enrichment score for the human PrLD and the mean SG enrichment score for its yeast PrLD counterpart.

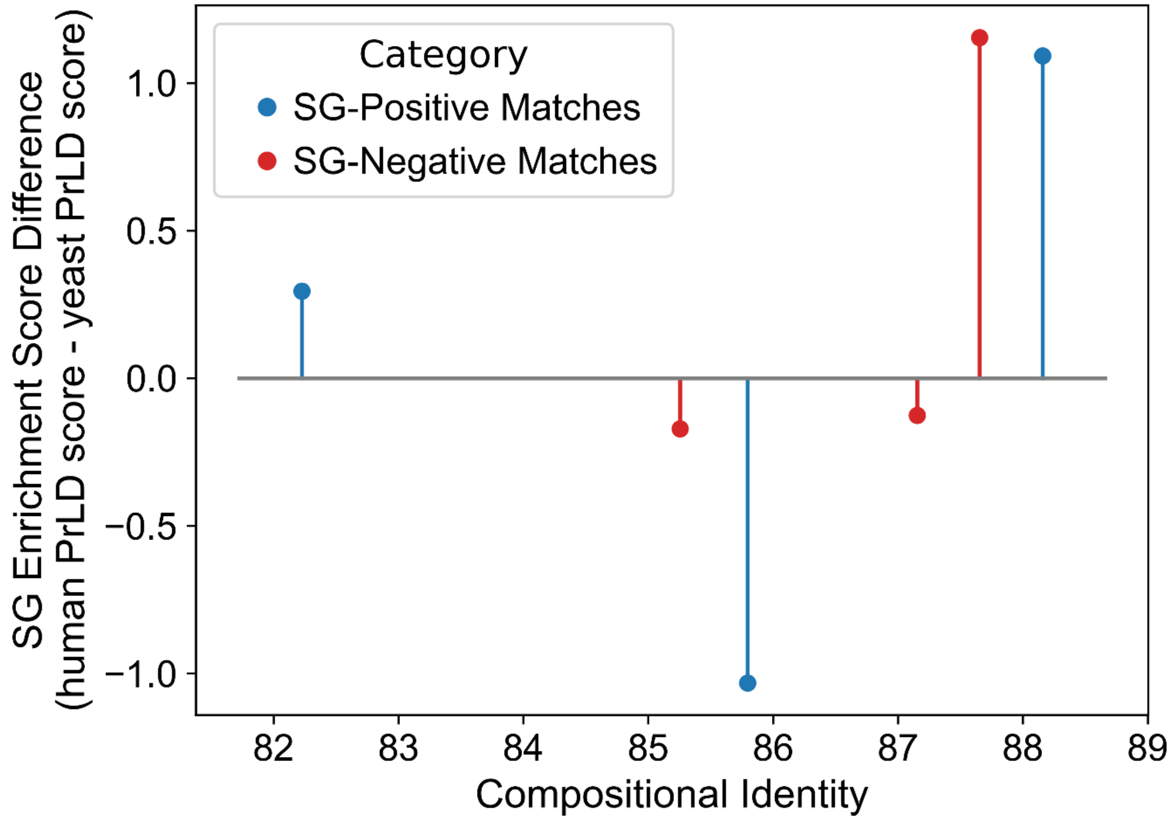

### Supplementary Table Legends

**Table S1. MatchIDR search results from *Saccharomyces cerevisiae* for the sPrLD and cPrLD query proteins.**

**Table S2. *p*-values for SG enrichment scores.** Two-tailed Welch's *t*-tests were performed to determine if the degree of SG localization differed significantly from a protein with no detectable SG enrichment (cPrLD2). For each comparison, the SG enrichment scores of that protein to the SG enrichment scores for cPrLD2 from Fig 2.

**Table S3. MatchIDR search results from *Homo sapiens* for the sPrLD and cPrLD query proteins.**

**Table S4. MatchIDR search results from *Dictyostelium discoideum* for the sPrLD and cPrLD query proteins.**

**Table S5. MatchIDR search results from *Homo sapiens* for the native yeast PrLD query proteins.**
